# Membrane Permeability Drives the Extreme Potency of Fentanyl but not Isotonitazene

**DOI:** 10.1101/2025.07.22.666146

**Authors:** Joseph Clayton, George J. Farmer, Jacqueline Glenn, Shailesh N. Mistry, J. Robert Lane, Lei Shi, Lidiya Stavitskaya, Meritxell Canals, Jana Shen

**Affiliations:** Division of Applied Regulatory Science, Office of Clinical Pharmacology, Office of Translational Sciences, Center for Drug Evaluation and Research, United States Food and Drug Administration, Silver Spring, MD 20993, U.S.A.; Department of Pharmaceutical Sciences, University of Maryland School of Pharmacy, Baltimore, MD 21201, U.S.A.; Division of Biomolecular Science and Medicinal Chemistry, School of Pharmacy, and University of Nottingham Biodiscovery Institute, University Park, Nottingham NG7 2UH, U.K.; Centre of Membrane Proteins and Receptors, Universities of Nottingham and Birmingham, Midlands NG7 2UH, U.K.; Division of Physiology, Pharmacology and Neuroscience, School of Life Sciences, University of Nottingham, Queens Medical Centre, Nottingham NG7 2UH, U.K.; Computational Chemistry and Molecular Biophysics Section, National Institutes on Drug Abuse - Intramural Research Program, Baltimore, MD 21224, U.S.A.

## Abstract

Fentanyl is a leading cause of drug overdose deaths in the United States, yet the mechanisms underlying its extreme in vivo potency remain poorly understood. Recently, new synthetic opioids nitazene derivatives have emerged, among which isotonitazene is 50 times more potent than fentanyl. Here we used state-of-the-art molecular dynamics (MD) simulations and experiments to investigate the membrane-dependent pharmacology of fentanyl, isotonitazene, morphine, and naloxone. Using the weighted-ensemble continuous constant pH MD, we estimated the effective permeability of fentanyl at pH 7.5 to be on the order of 10^−7^ cm/s, which is about two orders of magnitude faster than the simulation estimate for morphine. In contrast, isotonitazene and naloxone effectively do not partition into the membrane under the same conditions. The simulations captured the proton-coupled permeation processes, challenging as well as refining the long-standing pH-partition hypothesis. Subsequent BRET reporter cell experiments demonstrated that cells exposed to fentanyl, but not morphine, reactivated the receptor after washout and in competition with naloxone. Immobilized affinity membrane chromatography confirmed fentanyl’s high affinity for phospholipids. Our findings strongly support the hypothesis that fentanyl’s extreme in vivo potency may be driven by its accumulation within the plasma membrane or intracellularly, enabling it to repartition into the extracellular space to rebind the receptor, or potentially access it via a lipid-mediated route. This highlights the importance of membrane-dependent pharmacology for understanding opioid toxicity and guiding the design of more effective antagonists. Our simulation methodology enables accurate prediction and analysis of membrane permeation of ionizable molecules, providing a valuable tool for ADME optimization in drug development.

## Introduction

The opioid-related deaths in the United States have increased sharply over the last decade. A new height was reached in 2023, with over 80,000 overdose deaths, among which 90% involved fentanyl,^1^ which is an ultrapotent synthetic (UPS) opioid exhibiting distinct pharmacology compared to the natural opiate morphine (Fig. 1A). Comparison of cryogenic electron microscopy (cryo-EM) structures of fentanyl- and morphine-bound *μ*OR^2^ suggests that fentanyl’s increased potency may result from the additional interactions with the receptor, particularly between its phenylethyl group and a minor pocket in *μ*OR.^3^ These unique receptor interactions likely contribute to the ∼ 10-fold higher in vitro functional potency of fentanyl compared to morphine;^4^ however, they cannot explain the drastically greater analgesic potency of fentanyl, which is 50–400 times that of morphine.^5^ In contrast, recent experiments using whole-cell patch-clamp electrophysiology and signaling assays found that fentanyl, but not morphine, can reactivate *μ*OR after washout, implying membrane retention.^6^ This lends support to an alternative hypothesis: fentanyl partitions into the plasma membrane, forming a local “drug depot” that may enhance binding kinetics and/or enable lipid-mediated receptor access.^7^ The hypothesis is consistent with the exceptional lipophilicity of fentanyl,^5^ as reflected by its octanol-water partition coefficient, which is over 700 fold higher than that of morphine^8^ despite their similar solution p*K*_a_ values – a discrepancy that remains poorly understood.

**Figure 1.**
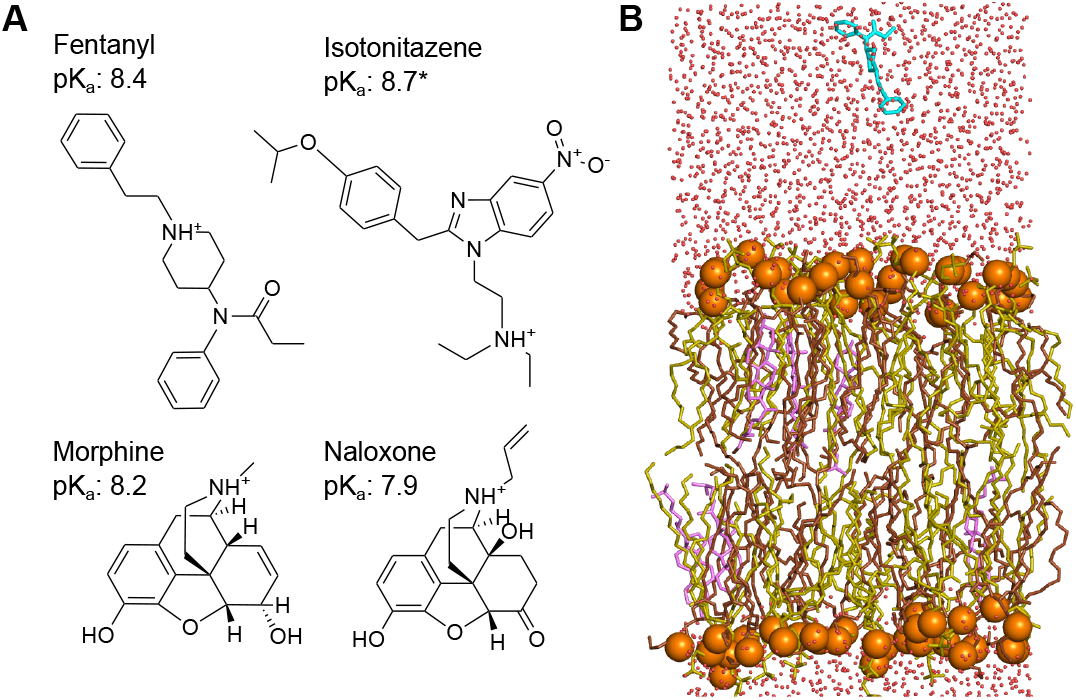
Chemical structures of fentanyl, morphine, isotonitazene, naloxone, and the simulation setup. **A**. Chemical structures and experimental solution p*K*_a_ values of fentanyl, ^12^ isotonitazene (approximated by that of dimethyltryptamine ^14^), morphine, ^13^ and naloxone. ^15^ **B**. A snapshot of fentanyl (cyan) approaching the model lipid bilayer comprised of POPC (yellow), POPE (brown), and cholesterol (pink) in a 5:5:1 ratio. This composition was chosen to mimic mammalian neural soma. ^25^ Water molecules are represented by red dots, and the lipid phosphorous atoms are shown as orange spheres.

Motivated by the unresolved questions surrounding fentanyl’s extreme potency and lipophilicity and to test the “drug depot” hypothesis, we set out to investigate the membrane permeation properties of three *μ*OR agonists (fentanyl, morphine, and isotonitazene) and the *μ*OR antagonist and opioid reversal agent naloxone (Fig. 1B) using state-of-the-art molecular dynamics (MD) simulations and experiments. Compared to fentanyl, isotonitazene is a newly emerged ultrapotent synthetic opioid from the 2-benzylbenzimidazole family; it was first detected around 2019 in the Midwest of the United States. Of note, isotonitazene is about 50 times more potent than fentanyl.^9^ Fatal overdose deaths involving isotonitazene prompted the Drug Enforcement Administration (DEA) to temporarily place it under Schedule I substances in 2020, with permanent scheduling in 2021.^10^ Isotonitazene has also raised significant concerns in Europe, contributing to a second wave of drug-related deaths in the UK.^11^ Fentanyl, morphine, and isotonitazene belong to distinct structural families (Fig. 1A), yet they share similarly basic solution p*K*_a_’s of 8.2– 8.7,^12–14^ which result in about 10% neutral species at physiological pH 7.5. Naloxone, in contrast, shares a similar structure with morphine but has a somewhat lower p*K*_a_ of 7.9.^15^

To allow direct simulation of membrane permeation processes of titratable molecules with full atomic detail, we integrated the GPU-accelerated particle-mesh Ewald continuous constant pH MD (CpHMD),^16,17^ which captures proton-coupled conformational dynamics, with the weightedensemble (WE) protocol,^18,19^ which accelerates sampling of rare events such as drug permeation.^20^ Traditional theoretical studies of passive membrane permeation by small molecules have relied on calculating free energy profiles (potential of mean force) along the membrane normal using umbrella sampling^21–23^ or biased sampling protocols such as metadynamics.^21,24^ In these methodologies, the permeant does not explore the full orientational degrees of freedom in the membrane environment due to limited sampling time in each umbrella window or the use of biasing potential in metadynamics. Here we report what we believe are the first atomic-level simulations to simultaneously depict full conformational flexibility, orientational freedom, and protonation state dynamics of drug molecules throughout the membrane permeation process. Additionally, the opioids examined here are significantly larger than the molecules studied by the previous simulation work.^20–24^ Findings from our simulations and experiments offer compelling support for the “drug depot” hypothesis and an atomically detailed explanation for fentanyl’s exceptional lipophilicity. More broadly, proton-coupled weighted-ensemble MD offers a powerful approach for investigating the membrane permeation of ionizable drugs—a common yet poorly understood aspect of drug pharmacokinetics.

## Results and Discussion

### Fentanyl partitions into the membrane orders of magnitude faster than other opioids

The membrane permeation simulations at 1 bar, 300 K, and solution pH 7.5 were conducted for three opioid agonists (fentanyl, morphine, and isotonitazene) and one antagonist (naloxone) using the PME-CpHMD module^17^ in Amber24.^26^ In order to accelerate the sampling of rare events (i.e., membrane permeation), the weighted ensemble method^18^ implemented in WESTPA 2.0^19^ was employed, with the progress coordinate defined as the permeant center of mass (COM) *z*-position relative to that of the lipid bilayer. The simulations were conducted under steady-state conditions and initiated with the permeant placed approximately 10 Å away from a fully solvated lipid bilayer composed of palmitoyloleoylphosphatidylcholine (POPC), palmitoyloleoylphosphatidylethanolamine (POPE) and cholesterol in a 5:5:1 ratio (Fig. 1B and SI Fig. S1). All molecules primarily adopt the charged (protonated amine) state in solution (SI Fig. S2). To confirm our findings, a second set of simulations was conducted for all molecules with a modified WE protocol, where additional WE bins were added near the upper leaflet-water interface to further enhance sampling of the membrane partitioning events. Thus, our discussion will focus on this second set of simulations; unless otherwise noted, results from the first set are given in parentheses.

The simulations demonstrate that the membrane permeation rate follows the order: fentanyl *>* morphine *>* isotonitazene *>* naloxone. Based on the probability flux,^20^ we estimated the mean first passage time (MFPT), which is the average time for the permeant to pass the membrane for the first time. The estimated MFPT of fentanyl is 10 (4.6) s, while those of morphine, isotonitazene, and naloxone are orders of magnitude larger, at 9.8 × 10^2^ (8.8 × 10^5^) s, 3.9 × 10^12^ (3.0 × 10^9^) s, and 1.8 × 10^30^ s, respectively (Table 1). Note, the MFPT of naloxone could not be estimated in the first trial, as it did not permeate the lower leaflet within the simulation time (SI Fig. S3). These data suggest that: 1) fentanyl is the most capable of partitioning into the membrane; 2) morphine can partition into the membrane, albeit much more slowly than fentanyl, and; 3) isotonitazene and naloxone are unable to partition on physiologically relevant timescales. Surprisingly, naloxone’s MFPT is orders of magnitude larger than morphine’s despite their highly similar structures.

**Table 1:** Effective permeability and mean-first passage time estimated from simulations and chromatographic hydrophobicity index estimated from experiment.

| | $\log(P_m[\text{cm s}^{-1}])$ | MFPT (s) | CHI |
| --- | --- | --- | --- |
| Fentanyl | -7.7 [-7.9,-7.5] | 10. [7.4,16] | 41 |
|  | -7.3 [-7.5,-7.2] | 4.6 [3.4,6.9] |  |
| Morphine | -9.6 [-9.9,-9.5] | 9.8e2 [6.2e2,1.8e3] | 17 |
|  | -12.6 [-12.7,-12.5] | 8.8e5 [7.1e5,1.2e6] |  |
| Isotonitazene | -19.2 [-19.7,-19.0] | 3.9e12 [2.3e12, 1.1e13] | n/d |
|  | -16.1 [-16.4,-15.9] | 3.0e9 [1.9e9,6.2e9] |  |
| Naloxone | -36.9 [-37.0,-36.8] | 1.8e30 [1.5e30,2.4e30] | 28 |

Based on the probability flux and an effective reaction volume,^20^ we also estimated the membrane permeability coefficient *P*_*m*_ [cm s^-1^] to compare with other drug molecules (Table 1). Fentanyl has an estimated log*P*_*m*_ around -7.5 (-7.3), which is much lower than the experimental log*P*_*m*_ of water (-4) but only one order of magnitude lower than the small, neutral drug-like compounds zacopride, sotalo, and tacrine (between -5 and -6).^20^ In contrast, the estimated log*P*_*m*_ values of morphine and isotonitazene, -9.7 (-12.6) and -19.2 (-16.1), respectively, are comparable to the experimental log*P*_*m*_ of potassium ions (-14),^27,28^ while the estimated log*P*_*m*_ of naloxone is -36.9, which is orders of magnitude lower than all the opioid agonists.

Note, the log*P*_*m*_ difference between the first and second trials for morphine and isotonitazene is roughly three orders of magnitude (Table 1), indicating large statistical uncertainties likely due to insufficient sampling of rare permeation events. However, the log*P*_*m*_ differences between morphine and isotonitazene – and in fact between any two opioids – are sufficiently large that their 95% intervals from both trials do not overlap (for naloxone, only one interval was estimated, Table 1). The statistical significance of the log*P*_*m*_ difference between fentanyl and morphine is further supported by a two-sided t-test using the last 50 WE iterations of the two trials, which yielded a *p*-value of 0.0001, well below the threshold of 0.05. Therefore, while individual permeability estimates have large uncertainties, the relative order among opioids is robust.

### Membrane permeability is enhanced by early deprotonation within the membrane

To understand the distinct membrane permeation behavior of the four molecules, we examined several microscopic quantities of the permeant as a function of its *z*-position, including the free energy profile (i.e., potential of mean force), fraction of deprotonation, change in the local membrane thickness, number of the first-solvation-shell water molecules, number of hydrogen bonds (Hbonds) and hydrophobic contacts (Fig. 2). The *z*dependent free energy (FE) profiles display a maximum at the membrane center for all molecules studied (Fig. 2C). Consistent with the trend in permeation rates, fentanyl exhibits the lowest FE barrier at 7.8 (11.2) kcal/mol, followed by morphine at 14.2 (18.8) kcal/mol, then isotonitazene at 24.8 (20.1) kcal/mol, and finally naloxone with the highest barrier at 33.4 (38.9) kcal/mol. To understand how these molecules that are charged in solution enter the hydrophobic membrane core, we examined the deprotonation profiles along *z* (Fig. 2D). Fentanyl remains protonated when its COM is close to the phosphate headgroups (*z* 20 ≈ Å) and begins deprotonation around 15 Å, becoming fully deprotonated below *z* ≈ 10 Å. Morphine begins deprotonation earlier, just below the phosphate headgroups, and becomes fully deprotonated slightly later than fentanyl, below *z* ≈ 7 Å. Despite sharing a highly similar structure with morphine, the titration profile of naloxone resembles that of isotonitazene, with early deprotonation onset but complete deprotonation only near the membrane center. These data demonstrate that the ability of the weakly basic permeant to complete deprotonation in the membrane is correlated with its permeability.

**Figure 2.**
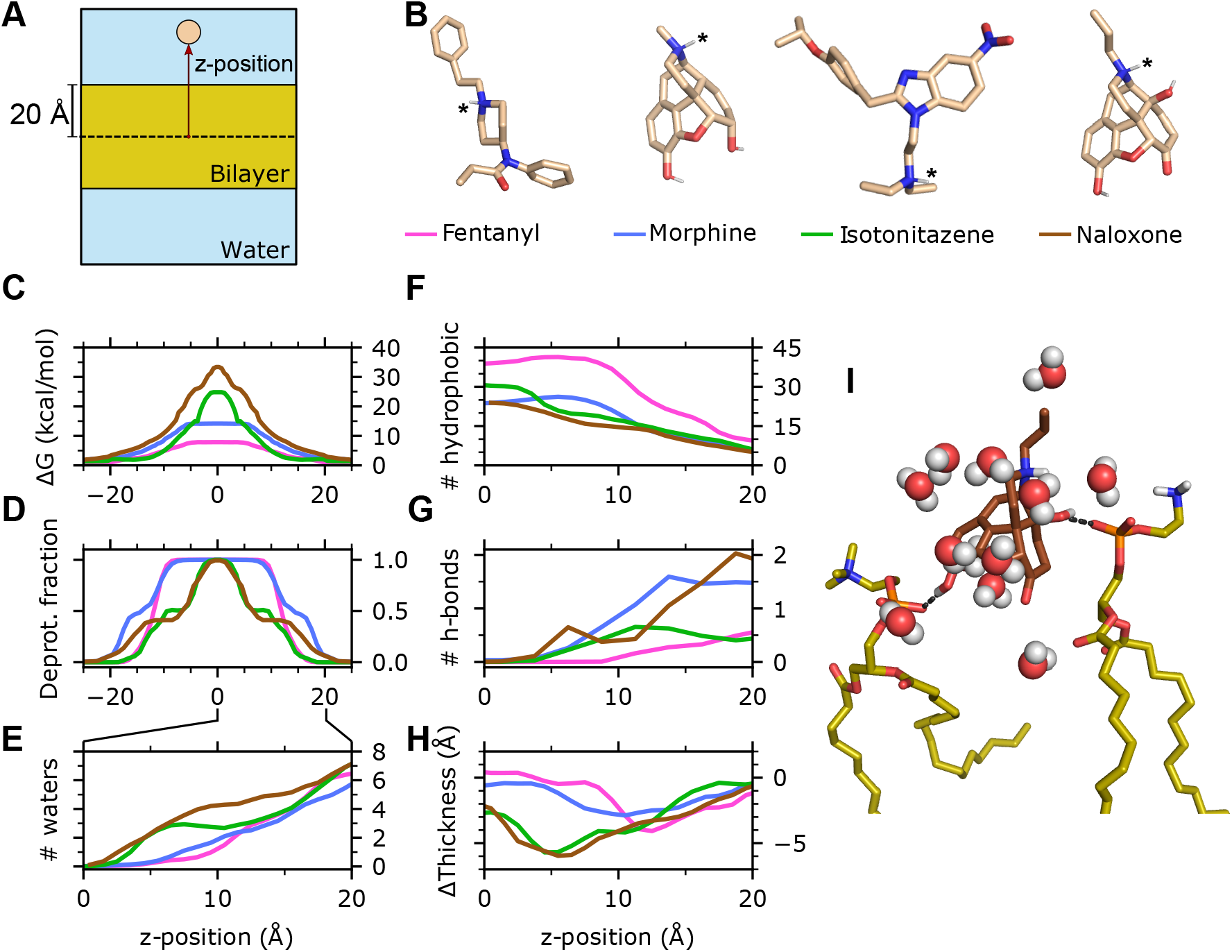
Molecular determinants of the distinct membrane permeability of fentanyl in comparison to morphine, isotonitazene, and naloxone. **A**. Illustration of the simulation system and *z* position of the permeant center of mass (COM). **B**. 3-D structures of the four compounds. Titratable nitrogen is indicated by an asterisk. Nonpolar hydrogens are hidden. The line color next to the permeant names corresponds to the data shown in **C-H. C**. Free energy profile (**C**) and deprotonation fraction (**D**) of the permeant along the permeant *z*-position. The profiles are symmetrized about *z* = 0 following Ref. ^20^ **E**. Average number of water molecules within 3.4 Å from any heavy atom of the permeant as a function of *z*. **F**,**G**. Number of hydrophobic contacts (**F**) and H-bonds (**G**) between the permeant and lipid molecules as a function of *z*. **H**. Change of the membrane local thickness around the permeant as a function of *z*. The local thickness is defined as the *z* distance between the centers of phosphorous atoms in the upper and lower leaflets with a 10-Å cylinder around the permeant COM. The average value of the local thickness when the permeant COM is *>* 30 Å from the membrane is used as a reference. Data for **C-H** are taken from the final 100 iterations of the second set of WE-CpHMD simulations (see SI Fig. S6) for the first set of simulations). **I**. A trajectory snapshot shows that naloxone (brown) forms two H-bonds between its hydroxyl groups and the phosphate groups of two lipids while its charged amine interacts with several water molecules.

### Hydrophobicity and diminished hydrogen bonding capacity drive the distinct titration-dependent permeation behavior of fentanyl

In solution, the charged amine of the four molecules is stabilized through hydrogen bonding (H-bonding) with water. During titration-coupled membrane partitioning, the number of water molecules surrounding the permeant decreases. Fentanyl and morphine become completely desolvated below *z* ≈ 5 Å, while isotonitazene and naloxone retain 1–2 water molecules until reaching closer to the bilayer center (Fig. 2E and SI Fig. S6). The earlier desolvation of fentanyl and morphine within the membrane is consistent with their earlier deprotonation compared to isotonitazene and naloxone. This raises the question, why does fentanyl permeate the membrane drastically faster compared to all other molecules? Important clues are provided by the hydrophobic and H-bond interaction profiles along *z* (Fig. 2F and G; SI Fig. S6), which correlate with the titration profiles. Throughout the membrane partitioning process, fentanyl forms the largest number of hydrophobic contacts in comparison to all other molecules, starting at about 10 and increasing to about 40 below *z* ≈ 10 Å, compared to 5–25 for morphine. This corroborates its 700fold higher P_ow_ relative to morphine.^8^ At the same time, fentanyl forms the least number of H-bonds with lipids, with an average occupancy of 0.5 at the level of the phosphate group and decreases to zero below *z* ≈ 10 Å. In contrast, the hydroxyl groups of morphine and naloxone are capable of forming H-bonds with the phosphate groups of two lipids at the membrane-water interface, and one H-bond persists until *z* approaches 5 Å (Fig. 2I).

### Interactions between the charged amine and water slows down permeation and causes local membrane distortion

Given the nearly identical structures and the similar number hydrophobic and H-bonding contacts, the significantly slower permeation rate of naloxone relative to morphine is puzzling. Simulations revealed that the delayed titration of naloxone is correlated with the persistent interactions between the charged amine and water molecules until approaching the bilayer center (Fig. 2D). Since these water molecules simultaneously interact with surrounding lipids (Fig. 2I), local lipids are displaced downward, reducing local membrane thickness. Interestingly, upon naloxone deprotonation, the membrane thickness recovers to normal values (Fig. 2H). The relationship between the hydration of the charged amine of naloxone and membrane thinning is corroborated by the first set of WE-CpHMD simulations, where the membrane exhibits progressive thinning as naloxone moves toward the bilayer center while preserving interactions between its charged amine and two water molecules (SI Fig. S6). The delayed titration of isotonitazene is also correlated with the persistent interactions with water and the resulting local membrane thinning (Fig. 2D,E,H). We suggest that the persistent interaction between the charged amine and water molecules is another significant contributor to the reduced permeation rates of naloxone and isotonitazene.

### Fentanyl inserts vertically at the membrane-water interface but adopts random orientations within the bilayer

To further understand the extraordinary membrane permeability of fentanyl, we analyzed its conformational dynamics during the permeation process. The orientation of fentanyl is defined using an angle formed between the membrane normal and a vector drawn from the amide nitrogen to the piperidine amine nitrogen (Fig. 3A). In solution, fentanyl samples random orientations (Fig. 3A–C and SI Fig. S7); as it initiates the partitioning process at the bilayer-water interface (*z* ∼ 20 Å), a vertical orientation relative to the membrane (angle around 30^°^ or 160^°^) is preferred, as demonstrated by the trajectory snapshots and the bimodal free energy profiles (Fig. 3A,B,D and SI Fig. S7). This vertical orientational preference may reflect the elongated molecular geometry of fentanyl, which exhibits greater compatibility with lipid organization than the globular ‘Tshaped’ structure of morphine or naloxone, thereby facilitating rapid membrane permeation. Note, while both sets of WE-CpHMD simulations favor vertical orientations for fentanyl, the second set shows a modest preference for the downward conformation, where the phenethyl group is oriented toward the membrane interface (Fig. 3A,B,D). Interestingly, once past the lipid head groups, fentanyl is free to adopt other orientations, and in the middle of the bilayer, no orientational preference is observed (Fig. 3B,E, SI Fig. S7, Supplemental Movie 1). This orientation freedom in the membrane core region is also observed for morphine, isotonitazene, and naloxone (Supplemental Movies 2, 3, and 4), which may be attributed to the absence of directional interactions with lipids (i.e., h-bonds as shown in Fig. 3G).

**Figure 3.**
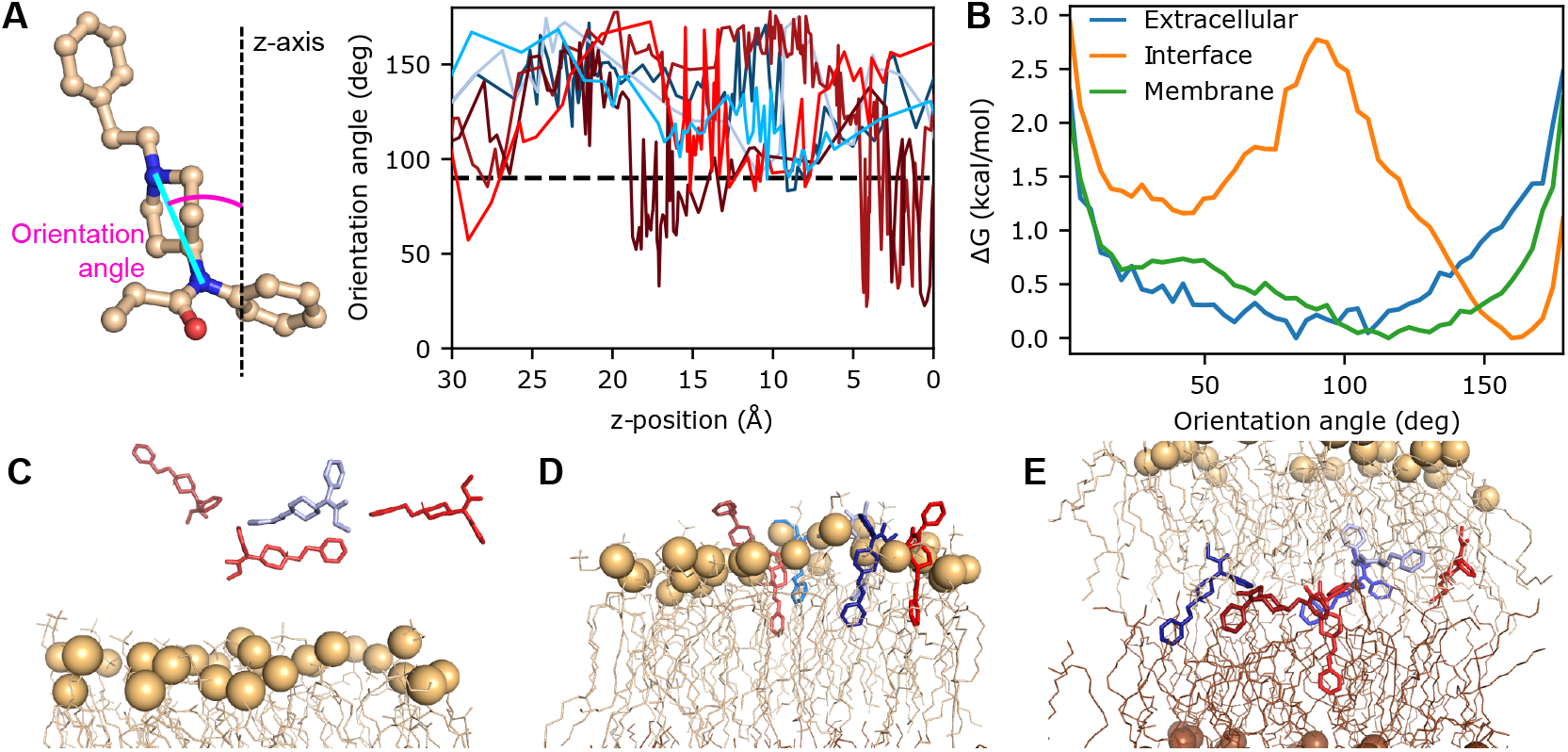
Fentanyl inserts vertically into the bilayer and adopts various orientations in the membrane. **A**. The orientational angle of fentanyl (left) as a function of *z* is calculated for unique continuous trajectories of fentanyl (right) permeating through the lipid bilayer. Angles greater than 90^°^ show the phenethyl group oriented downward. **B**. The free energy profile along the orientation angle while fentanyl is in three different regions: extracellular (blue; defined by *z*-position between 31 and 35 Å), at the upper leaflet interface (orange; *z*-position between 18 and 22 Å), and in the middle of the membrane (green; *z*-position between -2 and 2 Å). **C-E**. Snapshots selected from the seven trajectories, showing insertion into the upper leaflet (**C**, phosphorous atoms in tan), passage through the center of the bilayer (**D**), and insertion into the lower leaflet (**E**, phosphorous atoms in brown). Fentanyl is colored based on the trajectory the snapshot originated, matching **A**.

### Fentanyl is capable of reactivating the receptor after washout and even in competition with naloxone

To experimentally verify the unique membrane permeability of fentanyl, we developed a sensitive bioluminescence resonance energy transfer (BRET) protocol that harnesses the ability of *μ*OR to efficiently activate G_i_ G proteins upon binding of an opioid agonist. Note, experiments were not conducted for isotonitazene due to regulatory restriction on controlled substances. Cells expressing this BRET sensor were first stimulated with either 100 nM fentanyl or 1 *μ*M morphine. 10 minutes later, either 100 nM naloxone or the empty vehicle was injected. After another 15 minutes, three washouts were performed consecutively; these washouts were intended to completely remove any opioid agonist or naloxone even if it is initially bound to the receptor. The cells were then monitored for an additional 20 minutes, after which a high concentration of 10 *μ*M naloxone was applied.

As expected, naloxone injection caused a sharp decrease in BRET ratio for the fentanyl- or morphine-stimulated cells indicating that the receptor is largely inactivated, whereas vehicle injection (control) produced no change in BRET ratio (Fig. 4). Remarkably, following three washouts, the BRET ratios of the cells treated with fentanyl/vehicle (yellow) and fentanyl/naloxone (light brown) increase sharply, demonstrating that fentanyl reactivated the G protein complex even in competition with naloxone (Fig. 4). In contrast, minimal reactivation was shown by cells treated with morphine/vehicle (dark brown) or morphine/naloxone (purple). These observations suggest that fentanyl is retained by the cell membrane to a substantially greater degree than morphine, resulting in *μ*OR reactivation following washout.

**Figure 4.**
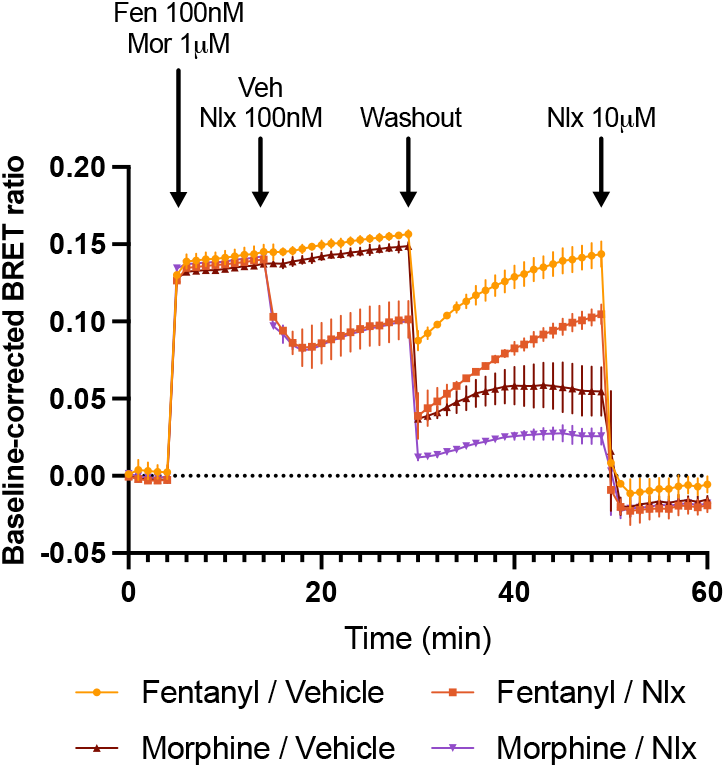
Fentanyl reactivated the *μ*OR after washout even in competition with naloxone. Cells were first transfected with *μ*OR and BRET G_i_ G protein activation sensor. After reading the BRET baseline, cells were stimulated with either 100 nM fentanyl or 1 *μ*M morphine. 10 minutes later 100 nM naloxone or vehicle was introduced, followed by three consecutive washouts. The baseline-corrected BRET ratios of cells stimulated with fentanyl/vehicle (yellow), fentanyl/naloxone (light brown), morphine/vehicle (dark brown), and morphine/naloxone (purple) were monitored for 20 minutes before finally introducing a high dose of 10 *μ*M naloxone.

### Fentanyl repartions into the extracellular solution after cell washout

To further test if cells can retain fentanyl, a separate experiment was conducted where non-transfected cells were exposed to a 10 *μ*M concentration of fentanyl, morphine, DAMGO, or naloxone for 30 minutes. The supernatant was then extracted, either immediately (No Wash) or after a series of 3-5 washes (Wash 3, Wash 4, and Wash 5). Reporter cells described above were then exposed to this supernatant (or in the case of naloxone, stimulated with 100 nM DAMGO) and the measured BRET ratio indicative of receptor activation or deactivation (in the case of naloxone) was compared to a direct exposure to a 10 *μ*M concentration of the corresponding opioid (Direct, Fig. 5).

**Figure 5.**
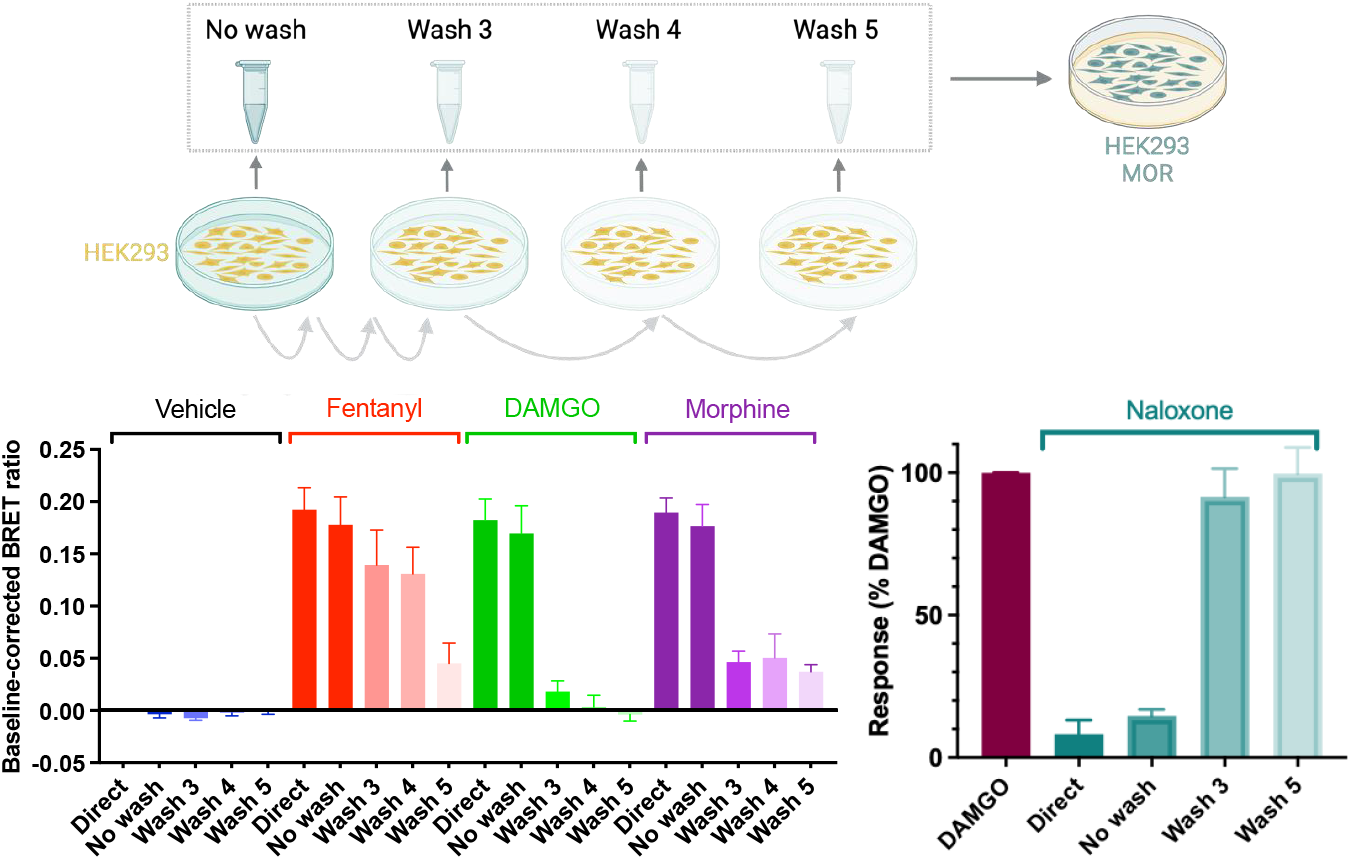
In contrast to morphine and naloxone, fentanyl repartitioned into the extracellular solution after cell washout. Untransfected cells were first incubated with a 10 *μ*M concentration of vehicle (blue), fentanyl (red), DAMGO (green), morphine (purple), or naloxone (turquoise) for 30 minutes; the supernatant of the cells was recovered either immediately (No Wash) or after a series of 3 to 5 washes (Wash 3-5). The supernatant was then added to separate cells transfected with *μ*OR and BRET G_i_ G protein activation sensor as in Fig. 4. In the case of naloxone, the transfected cells were first stimulated with 100 nM DAMGO (maroon). For comparison, the stimulation/inhibition from a direct application of the 10 *μ*M vehicle or opioid (Direct) was also measured.

Unsurprisingly, the transition from No Wash to Direct conditions produced only a slight decrease in agonist (fentanyl, DAMGO, or morphine)-induced receptor activation and antagonist (naloxone)-induced receptor deactivation (Fig. 5). Additionally it’s unsurprising that the response of DAMGO is significantly different after 3 washes compared to direct application, as DAMGO is a peptide and thus would not be able to permeate the cell membrane. The response of both morphine- and naloxone-incubated cells was also significantly weaker after 3 washes compared to direct application. Specifically, after 3 washes, receptor activation in morphine-incubated cells was reduced by about 80%, while that in naloxone-incubated cells was nearly restored to the level of DAMGO. In contrast, after 3 washes of cells exposed to fentanyl, a response similar to directly applied fentanyl remained (Fig. 5). In fact, for fentanyl a significant drop in activation was only observed in Wash 5.

Receptor activation by the supernatant following three washes of fentanyl-exposed cells suggests that fentanyl is retained intracellularly and subsequently repartitions into the extracellular solution. In contrast, morphine shows minimal cellular retention, while DAMGO and naloxone show nearly none. These data corroborate the WE-CpHMD simulation results showing fentanyl’s distinctively high membrane permeability, which exceeds morphine by two or more orders of magnitude and naloxone even more.

### Fentanyl exhibits greater retention in the immobilized artificial membrane compared to morphine and naloxone

Finally, to further test the membrane partition ability of the opioids, we conducted an experiment using Immobilized artificial membrane-high performance liquid chromatography (IAM-HPLC). The IAM column is comprised of POPC monolayers covalently attached to aminopropyl silica particles which are intended to mimic the in vivo interactions of drug molecules with phospholipids.^29,30^ Due to the monolayer construct, the analyte only binds to the phospholipid surface and does not pass through the membrane, in contrast to the simulation or physiological conditions where a bilayer exists.^30^ Nonetheless, the retention of the analyte in the column relative to the mobile phase, which is known as the chromatographic hydrophobicity index (CHI), provides a more accurate estimate of drug-membrane affinity compared to the octanol-water partition coefficient. IAM-HPLC yielded a CHI value of 41 for fentanyl, which is significantly higher than that for morphine (17) or naloxone (28, Table 1), indicating a higher affinity for phospholipids. This is consistent with fentanyl’s substantially higher simulated permeability and resistance to cell washout.

The higher CHI value of naloxone (28) relative to morphine (11) seems to contradict its lower simulated permeability and reduced resistance to cell washout. We suggest that the discrepancy may be primarily attributed to the aforementioned monolayer (IAM experiment) vs. bilayer construct (simulation and cell washout experiment). Second, morphine and naloxone share highly similar chemical structures; however, our simulations showed that naloxone forms more h-bonds with phosphate headgroups than morphine, prolongs its residence time in that region (Fig. 2G). This may explain the somewhat higher CHI index.

## Concluding Discussion

The WE-CpHMD simulations and experimental data suggest that fentanyl has an exceptional affinity for the cell membrane. The simulation-estimated effective *P*_*m*_ of fentanyl at pH 7.5 is on the order of 10^-7^ cm/s, which is at least two orders of magnitude larger than morphine. Our estimated permeability of fentanyl is 1–2 orders of magnitude smaller than nicotine, estimated as 10^-6^ cm/s at pH 7.4^31^ and 10^-5^ cm/s at pH 7.8^32^ by two independent studies. This difference is justifiable, considering that nicotine has a much smaller size but titrates at physiological pH with a similar p*K*_a_ value of 7.9.^32^ Note, compared with nicotine, similarly sized neutral drug-like compounds, zacopride, sotalo, and tacrine have similar permeability range of 10^-5^– 10^-6^ cm/s.^19^ Consistent with the simulation results, our experiments showed that fentanyl can reactivate the *μ*OR following washout and after naloxone displacement, whereas morphine cannot. These findings support the hypothesis that the membrane acts as a drug reservoir for fentanyl, either elevating local fentanyl concentrations or facilitating receptor binding through an alternative lipid-mediated pathway. This membrane-dependent mechanism may significantly contribute to the extreme potency of fentanyl relative to morphine. Note, fentanyl’s residence time at the receptor (about 3.8 minutes)^33,34^ is longer than morphine (about 0.72 mins);^34,35^ however, both residence times are significantly shorter than the cell’s exposure time (about 15 minutes) to the opioid before washout in our experiment. Therefore, the observed receptor reactivation by fentanyl after washout cannot be attributed to the longer residence time of fentanyl relative to morphine.

While most drugs are at least partially ionized in solution at physiological pH,^36,37^ the long-standing pH partition hypothesis posits that only the neutral form can traverse biological membranes.^38^ This hypothesis is supported by pH-dependent permeability profiles observed for ionizable drugs. For example, nicotine (p*K*_a_ values of 7.9)^32^ demonstrates increasing *P*_*m*_ with increasing pH,^31,32^ corresponding to a greater fraction of the neutral form.

In support of the pH partition hypothesis, our simulations demonstrated that only neutral species can cross the membrane; however, our simulations additionally revealed a more nuanced mechanism: weakly basic drugs can initially partition into the lipid environment in the protonated form and subsequently undergo deprotonation prior to reaching the bilayer center. This proton-coupled permeation mechanism is kinetically feasible because the timescale of deprotonation events, which is experimentally estimated at 1-10 *μ*s based on *k*_off_ on the order 1–10 x 10^5^ s^−1^,^39^ is orders of magnitude faster than that of membrane permeation events of most permeable drugs, which is estimated as 0.4 – 40 ms based on the effective permeability range of 10^−7^ – 10^−5^ cm/s at physiological pH^22,40^ and a membrane thickness of 40 Å.

To contextualize our findings, we compare our simulation results with previous theoretical studies of membrane permeation of small ionizable drugs using umbrella sampling PMF calculations based on either fixed-charge simulations^24,41^ or membrane-enabled hybrid-solvent CpHMD.^23^ Notably, fixed-charge simulations assume that ionized drugs can permeate the membrane, albeit at reduced rates, and thus the overall effective permeability at specific pH is determined by the fraction of the neutral species in solution.^23,24,41^ Our results align with the PMF calculations of propranolol (p*K*_a_ of 9.5) partitioning into the membrane using hybrid-solvent CpHMD simulations^23^ in demonstrating that ionizable drugs neutralize as they approach the hydrophobic membrane core. These observations are consistent with the experimental data^42^ and multi-site *λ*-dynamics simulations^43^ showing that ionizable residues in membrane-inserted peptides (with the exception of arginine) undergo large p*K*_a_ shifts that allow them to adopt the neutral state at physiological pH. Arginine maintains its positive charge, consequently inducing pore formation and membrane deformation;^42,44^ consistent with this, our simulations demonstrated that naloxone and isotonitazene, which remain partially protonated until reaching the membrane core, induce localized hydration and membrane thinning. Note, an important contrast between our results and the hybrid-solvent CpHMD umbrella sampling simulations of propranolol^23^ is that our calculated PMFs consistently exhibit energy barriers at the membrane center, which is in agreement with both fixed-charge umbrella sampling studies^24,41^ and the weighted ensemble permeation simulations of neutral molecules.^20^

The present simulations have several caveats.

While the relative permeability ranking among the four molecules is robust, the uncertainty in *P*_*m*_ values for poorly permeable molecules is substantial due to insufficient sampling of barrier-crossing events which are extremely rare. Notably, in the first set of simulations for naloxone, no permeating event was observed. Additionally, although WE simulations are theoretically unbiased, the selection of progress variables and other protocol parameters can influence the estimated *P*_*m*_ values, as demonstrated previously.^20^ Another limitation concerns the accuracy of force fields. Additive force fields such as CHARMM36 used in this work are known to overestimate the hydrophobicity of alkane environments such as the bilayer core, resulting in underestimated *P*_*m*_ values for polar molecules. For example, CHARMM36 underestimates water *P*_*m*_ in POPC bilayer by one order of magnitude.^45^ Given their ability to form h-bonds, the permeabilities of morphine, naloxone, and isotonitazene are likely underestimated. Nonetheless, the qualitative trend should remain valid as this systematic bias affects all opioids.

There are several caveats with the performed experiments. Whilst IAM-HPLC retention measurement more closely mimics the phospholipid membrane compared to the octanol-water partition estimates, it relies on POPC monolayers, which differ from the lipid bilayers of cell membranes. This may explain why it yielded a higher CHI value for naloxone relative to morphine, inconsistent with the simulation-estimated lower permeability and the observed increase in susceptibility to cell washout. The BRET washout experiment demonstrated that fentanyl is retained within cells and capable of reactivating the receptor; however, the precise mechanism underlying this reactivation remains unclear. Given fentanyl’s exceptional membrane permeability, it may accumulate within the lipid bilayer or intracellularly and repartition to the extracellular space, allowing it to rebind/reactivate the receptor. Alternatively, fentanyl may access the receptor binding pocket through a lipid pathway,^7^ which was recently investigated through mutagenesis of two residues located at the TM6/TM7 interface. However, these mutations did not alter fentanyl’s in vitro pharmacology.^6^

The present work offers compelling evidence to support the notation that plasma membrane accumulation is an important driver of fentanyl’s extreme potency, establishing a foundation for future studies to investigate the membrane-dependent mechanism of action of fentanyl and other opioids. Understanding membrane-dependent pharmacology has implications for designing new opioid antagonists. Recently, antagonists based on the fentanyl structure have been developed.^46^ Testing these antagonists in vitro and in vivo is a future direction of research towards mitigating the current opioid crisis. While regulatory restrictions prevented experimental validation with isotonitazene, our computational data demonstrate that it is unable to partition into the membrane at biological timescales. Instead, we hypothesize that isotonitazene’s potency is primarily driven by its high affinity for *μ*OR, as reflected by a subnanomolar *K*_*i*_ value approximately 12 fold lower than that of fentanyl.^47^ According to our recent computational study, nitro-containing nitazenes (e.g., isotonitazene) occupy subpocket 2 formed by TM1, TM2, and TM7, in contrast to fentanyl and morphine which engage subpocket 1 located between TM2 and TM3.^48^ This distinct binding mode of isotonitazene might also contribute to its enhanced potency relative to fentanyl.

## Materials and Methods

### WE-CpHMD simulations

#### CpHMD parameterization

The all-atom particle mesh Ewald continuous constant pH molecular dynamics (PME-CpHMD) method^16,17^ in Amber 2024^26^ was used for the membrane permeation simulations. In brief, each titratable site (either in a protein or a small molecule) is represented by a fictitious particle *λ* which is bound between 0 (protonated) and 1 (deprotonated) through an internal variable *θ, λ* = *sin*^2^ (*θ*), which is propagated simultaneously with the atomic coordinates. The CpHMD simulations necessitate two types of parameters: the p*K*_a_ of the model titratable site in solution (i.e. model compound) and the potential mean force (PMF) function of *λ* for the model compound. For fentanyl, morphine, and naloxone, the model p*K*_a_’s were 8.4,^12^ 8.2,^13^ and 7.9,^15^ respectively. As the p*K*_a_ of isotonitazene has not been experimentally determined, it was set to that of the structurally similar dimethytryptamine, 8.7.^14^ The parameters in the PMF functions were determined through thermodynamic integration (TI) simulations in water as described below. More details are given in the original CpHMD development work^16,17^ and a recent tutorial.^49^

The opioids were represented by the CGenFF force field.^50,51^ Water molecules were represented by the modified TIP3P model.^52,53^ The force field parameters of sodium and chloride ions were taken from Refs.^54,55^ For both TI and solution titration simulations for validation of the solution p*K*_a_ values of the opioids, a solvated system was built by solvating the opioid using a water box with a distance of at least 15 Å between the nearest water oxygen and the heavy atom on the small molecule. Sodium and chloride ions were added to reach an ionic strength of 0.15 M, with one additional chloride ion added to compensate for the net charge of 1 at pH 7.5. During the CpHMD titration, the effect of net charge was accounted for using a uniform background charge (plasma) in the PME correction term for propagating atomic coordinates.^16,17,56^ Each system was minimized for 5,000 steps, then seven independent replicas were created by fixing *θ*_*i*_ (0.2, 0.4, 0.6, 0.7854, 1.0, 1.2, or 1.4). Each replica was heated to 300 K under constant volume over 50 ps with a 5 kcal/mol/Å restraint on the heavy opioid atoms, then restraints were gradually relaxed over 100 ps under constant pressure of 1 bar. Pressure was controlled using the Monte-carlo barostat^57^ while temperature was controlled at 300 K using Langevin dynamics with a collision frequency of 1 ps^−1^. Long-range electrostatic interactions were calculated using particle mesh Ewald (PME) method with a real-space cutoff of 12 Å and a 1 Å grid spacing for the reciprocal space calculations. Lennard-Jones energies and forces were smoothly switched off over the range of 10 to 12 Å. The mean force ⟨*∂U* /*∂θ* ⟩ was then calculated over a 10-ns simulation for each *θ*; fitting the calculated mean force at a series of *θ* values to the partial derivative of the quadratic PMF function yields the two parameters in the PMF.^49^ Note that, in order to minimize fitting errors, fitting was done in the *θ* space and not the transformed *λ*.^49^ Following the TI simulations, titration simulations were performed to verify the experimental p*K*_a_’s in solution are recapitulated by CpHMD simulations. Six separate 20 ns simulations were conducted at different pH conditions, ranging from 7.0 to 9.5. To verify the parameters obtained from TI, the average deprotonation fraction over the final 10 ns of each simulation was measured then used to fit to the generalized Henderson-Hasselbach equation to obtain a calculated p*K*_a_(SI Fig. S4).

### System preparation for the WE-CpHMD simulations

To prepare the initial system, CHARMM-GUI^58^ was used to create a bilayer with a 5:5:1 ratio of POPC, POPE, and cholesterol; this composition was chosen to mimic mammalian neural soma^25^ and led to 66 lipids total (33 lipids per leaflet). An opioid was then induced and placed 30 Å from the center of the bilayer. A water layer of 22.5 Å was added above and below the bilayer. Sodium and chloride ions were added to reach an ionic strength of 0.15 M, and an additional chloride ion was added to neutralize the system at pH 7.5. The system was minimized for 5000 steps, then heated under constant volume to 300 K over 125 ps with restraints on lipid positions and dihedrals. These restraints were then gradually removed over 2.25 ns under constant pressure of 1 bar, followed by a 50 ns simulation to equilibrate the membrane. All steps were done using Amber24.^26^ The CHARMM36 lipid force field^59^ was used. All other parameters and settings were identical to those described above for the solution CpHMD simulations of opioids.

### The weighted-ensemble (WE) protocol

WE simulations, in brief, involve iteratively evolving a number of independent replicate systems called walkers. After each iteration a progress coordinate (or several coordinates) is calculated and used to place each walker into predefined bins; within each occupied bin, walkers are either replicated or removed such that the number of walkers per bin meets a target value. By updating a statistical weight for each walker after each split (replication) or merge (removal) both equilibrium and kinetic information can be estimated.

The progress coordinate used was defined as the z-position of the center of mass of the opioid (excluding hydrogen atoms), with the center of the membrane (defined by the center of all phosphorous atoms of POPC and POPE lipids) as the origin. Initial bin boundaries were set based on the environment: when the opioid was in solvent (|*z*| *>* 20 Å) boundaries were set 5 Å apart, while when inside the membrane (− 20Å *< z <* 20Å) boundaries were set 0.5 Å apart (85 bins total). This was chosen as diffusion within the membrane is likely slower than in the solvent. The WE protocol allows for dynamic bin boundaries, thus bins were added as needed at the membrane-extracellular interface (i.e. 20 *< z <* 25) in order to sample permeation. A steady state WE simulation was prepared, where a walker would be recycled if *z >* 55 Å or *z <* −25 Å at the end of the iteration. This was done to ensure the opioid does not cross the periodic boundary and that membrane partitioning occurs in one direction.

The WE simulations were conducted using WESTPA 2.0^19^ and Amber24.^26^ The target count for each bin was set to 4; each iteration involved simulating each walker for 100 ps at pH 7.5 using all-atom PME-CpHMD.^17^ WE simulations were conducted until both the calculated PMF along the progress coordinate and effective permeability had converged. All analysis was done using the WESTPA package and MDTraj.^60^ The rate of permeation was analyzed by dividing the system into three region based on the z-position from the bilayer center: extracellular (*z >* 20 Å), bilayer (−20 *< z <* 20 Å), and intracellular (*z <* −20 Å). The probability flux from extracellular to intracellular *f*_*ex → in*_ was measured using the w assign and w_direct tools within the WESTPA package,19,61 and then was used to calculate the effective permeability and mean first passage time (MFPT) following Zhang et al.:^20^

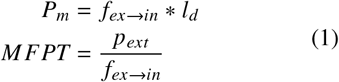

In the above equations, *l*_*d*_ represents the depth of the effective reaction volume while *p*_*ext*_ represents the fraction of trajectories that most recently sampled the extracellular state versus the intracellular state. The effective reaction volume is the region where the surface of the membrane influences neighboring molecules, thus events in bulk solvent (such as stirring) do not affect molecules in this region; following Zhang et al.,^20^ we set this to half of the height of the overall water buffer (*l*_*d*_ = 22.5 Å) in our analysis. Since the intracellular region was set as a recycle condition, there are no trajectories that sample the intracellular before sampling the extracellular–thus *p*_*ext*_ = 1. Statistical significance was determined by conducting a two-sided student’s t-test using the *f*_*ex → in*_ from each of the final 50 iterations.

The rate of deprotonation within the bilayer was estimated in a similar manner as described above. First, fentanyl was divided along two dimensions: the progress coordinate z-position (to determine if fentanyl was within the bilayer) and *λ* (to determine whether the amine is protonated or deprotonated). The bilayer region was defined as described above; the protonated state was defined as *λ <* 0.2 and the deprotonated state as *λ >* 0.8.

### Bioluminescence resonance energy transfer (BRET) experiments

#### Materials

DMEM was from Sigma-Aldrich, FBS was from Sigma-Aldrich, PEI was from PolySciences Inc, poly-D-lysine was from Fisher Scientific, coelenterazine h was from NanoLight Technology (Prolume Ltd), morphine was from Tocris, fentanyl was from Sigma-Aldrich, DAMGO was from Hello Bio, Naloxone was from Hello Bio and D-PBS was from Gibco.

#### Cell culture and transfection

Human embryonic kidney 293 T (HEK 293T) cells were cultured at 37 °C, 5% CO_2_ in Dulbecco’s modified eagle medium (DMEM) supplemented with 10% (v/v) fetal bovine serum (FBS). For transfection, cells were plated in 10 cm Petridishes (3 x 106 cells per dish) and allowed to grow overnight in full media at 37 °C, 5% CO_2_. 24h later, cells were transiently transfected, using a 1:6 total DNA to PEI ratio and the following DNA constructs: 2 *μ*M of G*α*i2, 1 *μ*M of G*β*1-Venus(156-239), 1 *μ*M of G*γ*2-Venus(1-155), 1 *μ*M of masGRK3ct-Rluc8 and 1 *μ*M of MOR [SNAPmMOR]. DNA/PEI mixtures were added to the cells. 24 post-transfection, cells were plated in Greiner poly-D-lysine-coated, white bottom 96-well plates (SLS) in full media.

#### BRET measurements

On the day of the assay (48h post-transfection), cells were washed once with D-PBS (Lonza, SLS) and incubated in D-PBS for 30 min at 37 °C. The Rluc substrate coelenterazine h was added to each well (final concentration of 5 *μ*M) and treated as specified below. BRET measurements (Venus and Rluc emission signals at 535 and 475 nm, respectively), were performed using a PHERAstar FSX microplate reader (BMG Labtech) at 37 °C. BRET ratio was calculated as the emission at 535 nm divided by the emission intensity at 475 nm signal and corrected for the vehicle BRET ratio signal. *Kinetic BRET experiments*. After a 5 min baseline read after the addition of coelenterazine h (final concentration of 5 *μ*M), morphine (1 *μ*M), fentanyl (100 nM) or vehicle were added, and the signal read for 10 min. Then, 100 nM of naloxone or vehicle were injected to the wells. At this concentration, naloxone was demonstrated to partially reverse the G protein activation evoked by morphine (1 *μ*M) and fentanyl (100 nM) to similar levels. The BRET signal was read for further 15 min prior to 3 washouts of the cells with D-PBS, re-addition of coelenterazine h (final concentration of 5 *μ*M) in drug-free D-PBS and measurement of the BRET signal for further 20 min until the final addition of 10 µM naloxone to fully reverse MOR induced G protein activation.

##### Ligand-release BRET sensor experiments

Untransfected HEK293T cells were incubated with vehicle or 10 *μ*M of fentanyl, morphine, or naloxone for 30 min at 37°C. After incubation, the supernatant of the cells was recovered (no wash) and cells washed, 5x with D-PBS and supernatants recovered (wash 1, 2,3,4, 5, respectively). 100 *μ*L of these supernatants were then used to stimulate a plate containing cells transfected with *μ*OR, and the G protein activation sensor constructs (G*α*i2, G*β*1-Venus(156-239), G*γ*2-Venus(1-155) and masGRK3ct-Rluc8). Direct application of vehicle or 10 *μ*M of fentanyl, DAMGO, morphine, or naloxone was used as a control. BRET signal was measured as above after 10 min incubation.

### IAM chromatography experiment

#### Equipment and methods

Immobilized artificial membrane-high performance liquid chromatography (IAM-HPLC) was performed using a conventional HPLC set up fitted with a IAM P.C DD2 column (30 x 4.6 mm, 10 *μ*M, 300 Å) (Regis technologies Inc, Chicago, USA). The column was maintained at 30°C with a flow rate of 1.5 mL/min and UV detection at 254 nm. The system consisted of a Shimadzu systems controller SCL-40, degassing unit DGU-405, solvent delivery module LC-40D XR, auto sampler SIL-40C XR, column oven CTO-40C and a photo diode array (PDA) detector SPD-M40 (Shimadzu, Kyoto, JPN). Solvent A contained ammonium acetate (50 mM) (SigmaAldrich, Gillingham, UK) in Milli-Q water at pH 7.4 and solvent B was acetonitrile (Thermo-Fisher Scientific, Loughborough, UK).

*Method 1*: 0-85% B over 4.75 min; 85% B for 2 min; 85–0% B over 0.5 min, then 0% B. All standards and samples were dissolved via dropwise addition of DMSO, before being diluted in water to 1 mM final concentration. Each sample was loaded in 10 *μ*L injections and repeated in triplicate on 2 separate days. IAM calibration mixture (Bio Mimetic Chromatography Ltd, Stevenage, UK) composition: octanophenone, heptanophenone, hexanophenone, valerophenone, butyrophenone, propiophenone, acetophenone, acetanilide, paracetamol. Test mixture 1: propranolol, indomethacin and colchicine; Test mixture 2: warfarin, carbamazepine, nicardipine. Samples: fentanyl, buprenorphine, naloxone, [D-Ala^2^, N-MePhe^4^, Gly-ol]-enkephalin (DAMGO), morphine all commercially available from standard suppliers.

#### Calibration plot

The system was calibrated by introducing the IAM calibration mixture (10 *μ*L) using method 1 and plotting the *t*_*R*_ of each component in the mixture against chromatographic hydrophobicity index (CHI (IAM)) values^62^ from the literature. Typical chromatograms, retention times (*t*_*R*_) and values of CHI (IAM) from the literature are shown in SI Figure S9. A calibration plot and equation for the line of best fit was generated in Microsoft Excel (version 16.88) and the Pearson correlation coefficient (r) was calculated to be *>*0.99, as summarized in graphical plot (SI Figure S11).

#### Column performance and suitability test

Before samples were analyzed, an assessment of column performance was carried out daily by introducing using Test mixtures 1 and 2 (SI Figure S11) using Method 1. The retention time of the components in each mixture were converted to CHI(IAM) values using the calibration plot in (SI Figure S10). The column was deemed suitable for analysis if all measured CHI (IAM) values were within ± 5 of their corresponding literature values^63^ Examples of typical chromatograms are shown in (SI Figure S11) and all data is summarized in SI Table S4.

#### Sample testing

Samples were analyzed using method 1 and typical chromatograms are shown in SI Figure S12. Retention time was converted to CHI (IAM) values for each sample using the calibration plot (SI Figure S10).

## Disclaimer

This article reflects the views of the authors and should not be construed to represent FDA’s views or policies. The mention of commercial products, their sources, or their use in connection with material reported herein is not to be construed as either an actual or implied endorsement of such products by the FDA. The contributions of the NIH author(s) are considered Works of the United States Government. The findings and conclusions presented in this paper are those of the author(s) and do not necessarily reflect the views of the NIH or the U.S. Department of Health and Human Services.

## Acknowledgments

J. C. was supported by the ORISE fellowship, which is a Research Participation Program at the FDA administered through the Oak Ridge Institute for Science and Education (ORISE) under the agreement between the FDA and Department of Energy. M. C. and R. L. were supported by the Academy of Medical Sciences (M. C.), Biotechnology and Biological Sciences Research Council (M. C., BB/T013966/1). G. F. was funded by the Welcome Trust Doctoral Training Programme in Drug Discovery and Team Science. J. S. was supported by the National Institutes of Health (R35GM148261). This research was supported in part by the Intramural Research Program of the National Institutes of Health (NIH), Z1ADA000606 (L. Shi).

## Supporting information

Supplemental information: supplemental tables and figures detailing simulation parameters, additional results from simulations, and IAM calibration/raw data. Supplemental movies : continuous trajectories showing the membrane permeation processes of fentanyl (movie 1); morphine (movie 2); isotonitazene (movie 3); and naloxone (movie 4).

## Author contributions

J. C.: conducted simulations and analyzed data, wrote and revised the manuscript; G.J. F: designed and performed IAM experiments and revised the manuscript; J. G.: performed BRET assays; S.N. M: designed and performed IAM experiments and revised the manuscript; M. C., R. L.: designed the project, supervised experiments, wrote and revised the manuscript; L. Shi. and L. S.: designed the project and revised the manuscript; J. S.: designed the project, analyzed the data, wrote and revised the manuscript.

## Data and materials availability

All simulation inputs and configurations can be found at https://github.com/JanaShenLab/Fentanyl-insert.

## TOC Graphic

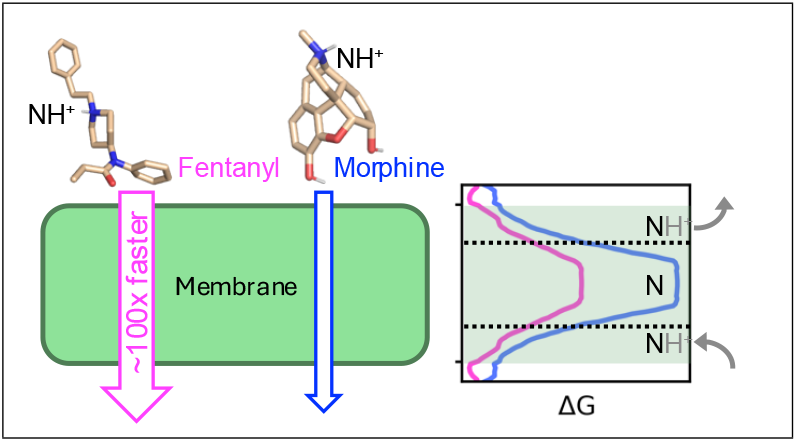

