## Supporting Information for "Membrane Permeability Drives the Extreme Potency of Fentanyl but not Isotonitazene"

### Supplemental Tables

Table S1: Parameters in the model titration PMF for the four compounds

| Molecule | A | B |
|---|---|---|
| Fentanyl | -58.059565 | 0.100857 |
| Morphine | -37.405443 | 0.136572 |
| Isotonitazene | -51.327454 | 0.325572 |
| Naloxone | -74.288134 | 0.046556 |

Table S2: Number of iterations and cumulative sampling time for each set of WE-CpHMD simulations

| Molecule | First trial | | Second trial | |
|---|---|---|---|---|
| | Iteration | Time | Iteration | Time ($\mu$s) |
| Fentanyl | 550 | 14.9 | 500 | 15.7 |
| Morphine | 550 | 11.3 | 500 | 15.5 |
| Isotonitazene | 400 | 8.5 | 350 | 9.5 |
| Naloxone | 300 | 3.4 | 350 | 7.0 |

Table S3: Physicochemical properties of four studied compounds

| Molecule | Mol weight | Rot. bonds | H-bond donor | H-bond acceptor | xLogP |
|---|---|---|---|---|---|
| Fentanyl | 336 | 6 | 0 | 3 | 3.94 |
| Isotonitazene | 410 | 9 | 0 | 4 | 3.62 |
| Morphine | 285 | 0 | 2 | 2 | 0.49 |
| Naloxone | 327 | 2 | 3 | 2 | 0.31 |

Molecular weight is unit of g/mol. xlogP (calculated octanol-water partition coefficient)[S1] values are taken from the Database IUPAC/BPS Guide to Pharmacology[S2] which reports the CDK calculations.[S3]

Table S4: Hydrophobicity index from immobilized affinity membrane (IAM) experiments

| Name | Acid/base profile | $t_R\pm$ SD (mins,n=6) | Measured CHI (IAM) | Reference CHI (IAM) |
|---|---|---|---|---|
| Propanolol | basic | 3.30 $\pm$ 0.04 | 47 | 42 |
| Nicardipine | basic | 3.31 $\pm$ 0.03 | 47 | 45 |
| Indomethacin | acidic | 2.13 $\pm$ 0.06 | 27 | 30 |
| Warfarin | acidic | 1.56 $\pm$ 0.06 | 18 | 20 |
| Carbamazepine | neutral | 2.14 $\pm$ 0.02 | 27 | 27 |
| Colchicine | neutral | 1.80 $\pm$ 0.07 | 22 | 23 |

Components of suitability Test Mixtures 1 and 2 and their corresponding acid/base profile. The average $t_R\pm$SD from n=6 repeats measured using method 1 was used to determine CHI(IAM) for each molecule using the calibration plot (Figure S10). The column was deemed suitable for analysis due to measured CHI(IAM) values being within $\pm$5 of reference CHI(IAM) from the literature.

#### Supplemental Figures

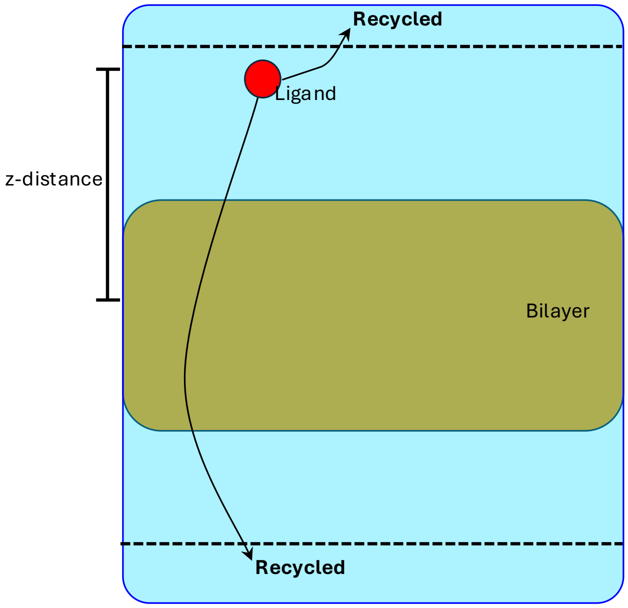


Figure S1: **Schematic of the steady-state condition.** Schematic of the steady-state condition used for the WE-CpHMD simulation. Once the ligand either moves far away from the membrane or permeates through, the walker is recycled–meaning the walker is terminated and a new one is started from the initial state.

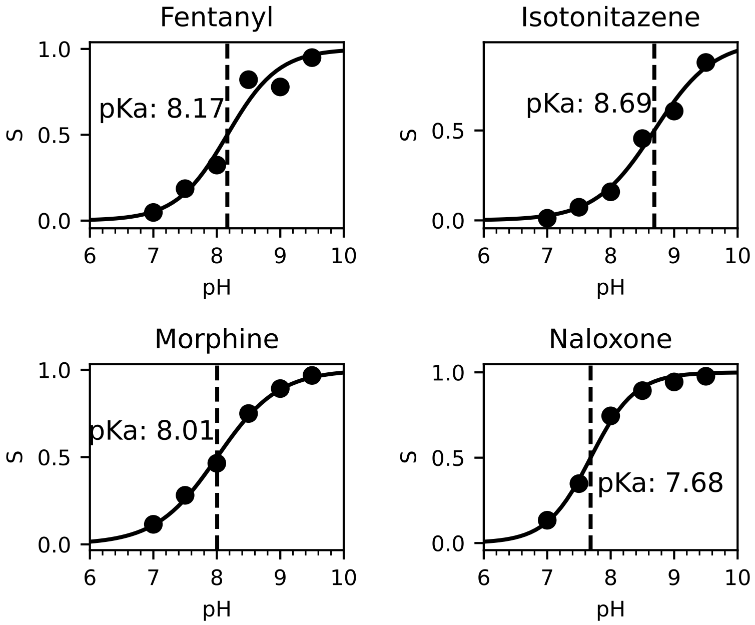


Figure S2: **Simulated solution titration curves of the four studied compounds.** Deprotonation fraction at different simulation pH conditions. 20 ns simulations were conducted at six different pH conditions ranging from 7.0 to 9.5. The final 10 ns of each was then used to calculate the deprotonation fraction. The calculated p$K_a$'s are given.

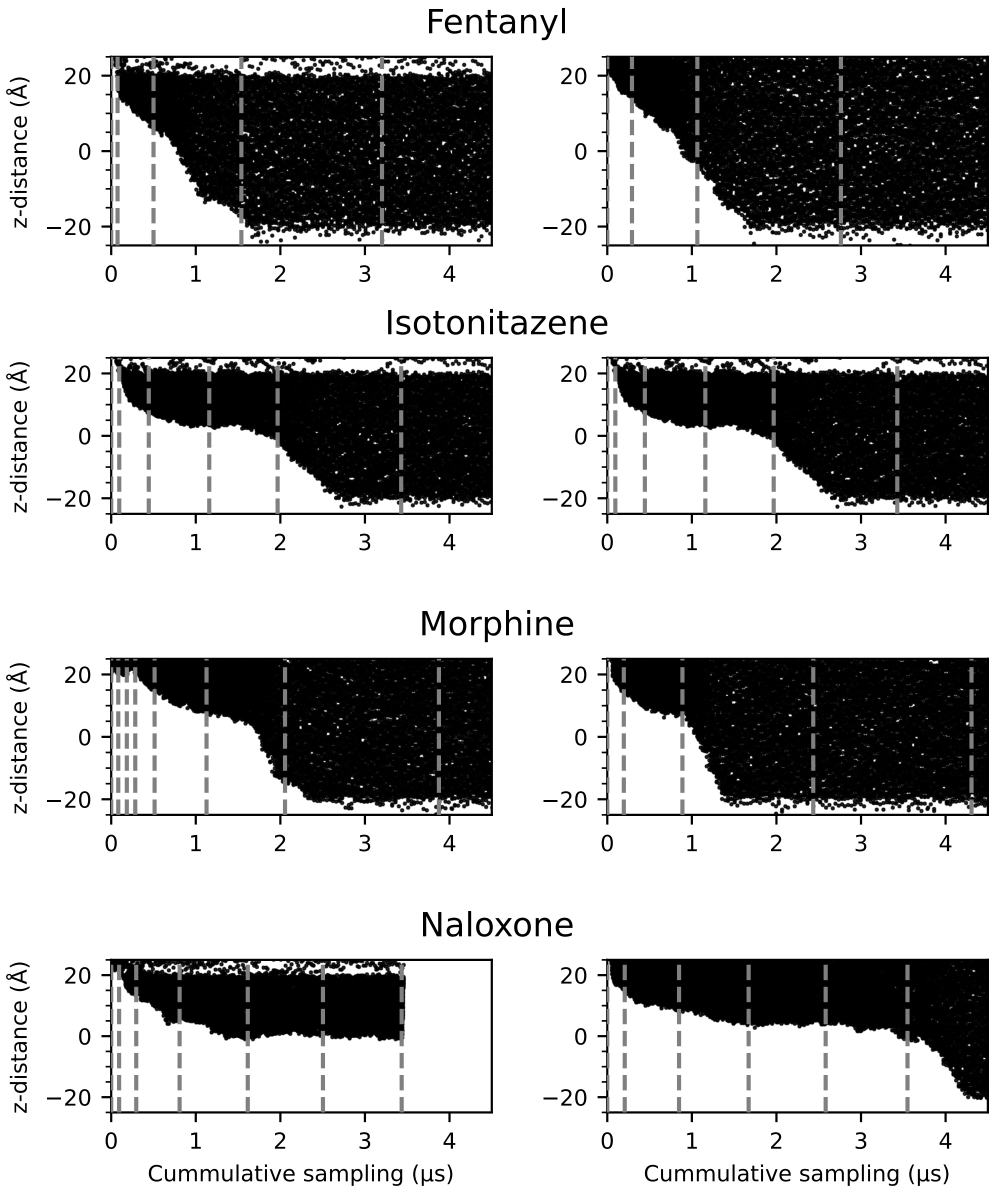


Figure S3: **Fentanyl requires the least simulation time to permeate the membrane.** $z$-position vs. cumulative simulation time of fentanyl, morphine, isotonitazene, and naloxone from the first (left) and second (right) sets of WE-CpHMD simulations. Gray dashed lines are drawn at every 50 WE iterations; for simplicity, only the first 4.5$\mu$s is shown.

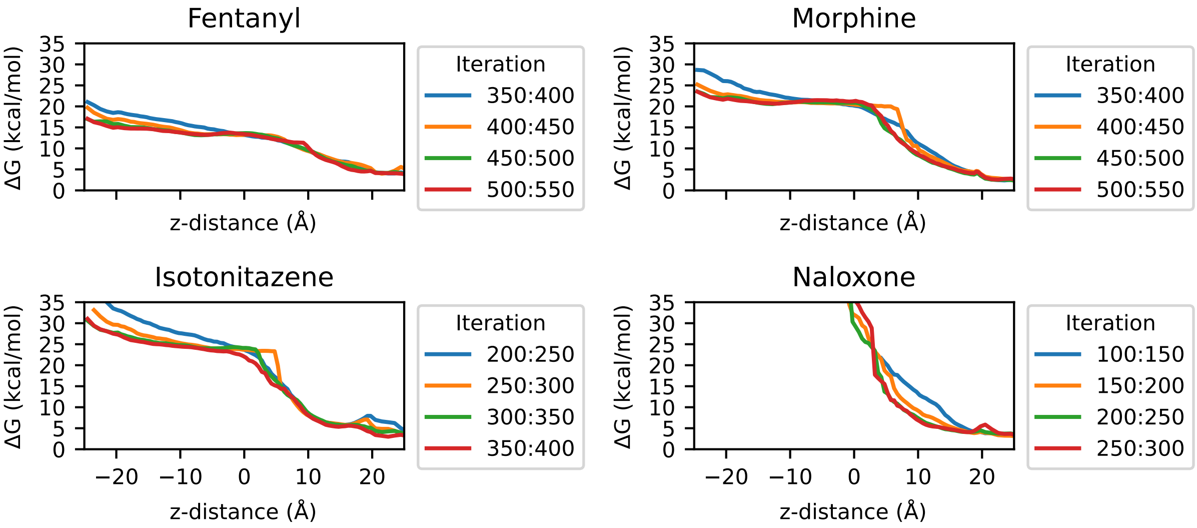


Figure S4: **Convergence of the potential of mean force calculations (first trial).** Convergence of the potential mean force, calculated using four windows containing 50 iterations, for the first set of weighted ensemble simulations.

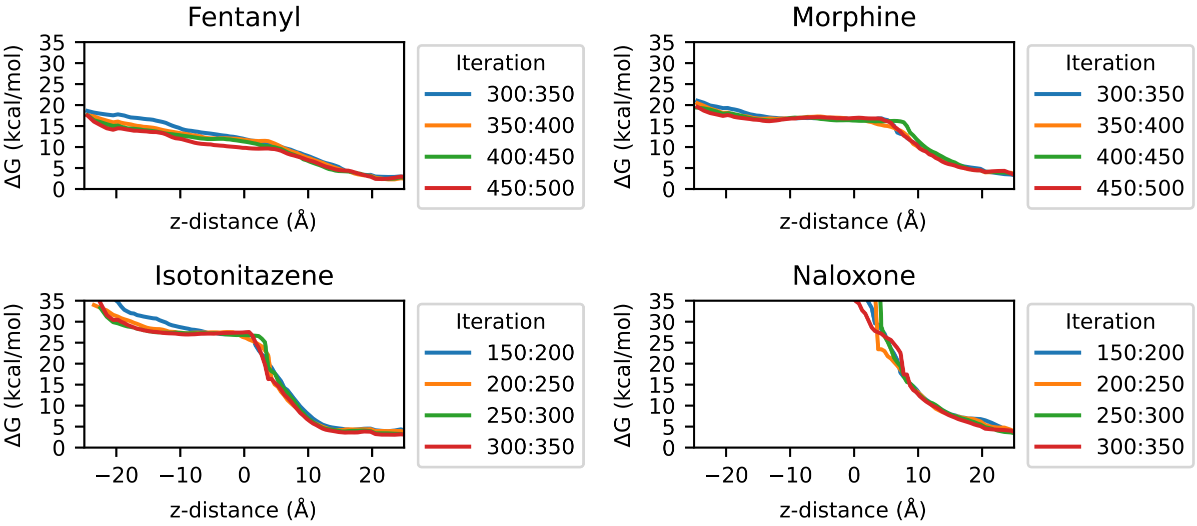


Figure S5: **Convergence of the potential of mean force calculations (second trial).** Convergence of the potential mean force, calculated using four windows containing 50 iterations, for the first set of weighted ensemble simulations.

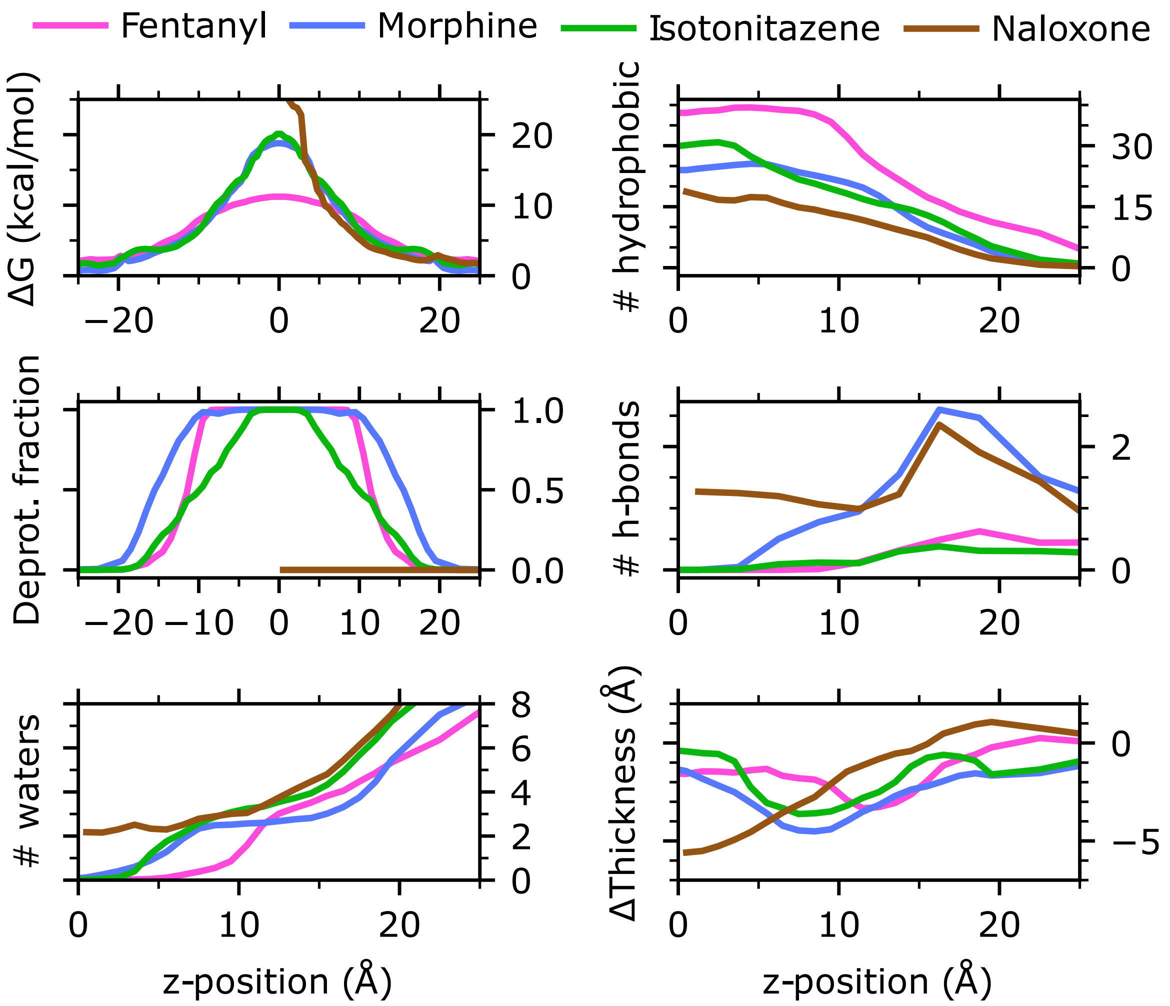


Figure S6: **Trends in permeation events are reproducible.** Characterization of the membrane permeation properties of fentanyl, morphine, isotonitazene, and naloxone from the simulation set 1. **Left.** Free energy profile (top), deprotonation fraction (middle) and the number of water within 3.4 Å from any heavy atom of the permeant (bottom) as a function of its $z$-position. The profiles are symmetrized about $z = 0$ following Ref.[S4] **Right.** Number of hydrophobic contacts (top) and h-bonds (middle) between the permeant and lipid molecules and the change of the membrane local thickness around the permeant (bottom) as a function of $z$. cylinder around the permeant COM. The average value of the local thickness when the permeant COM is $> 30$ Å from the membrane is used as a reference. Data are taken from the final 100 iterations of the first set of WE-CpHMD simulations.

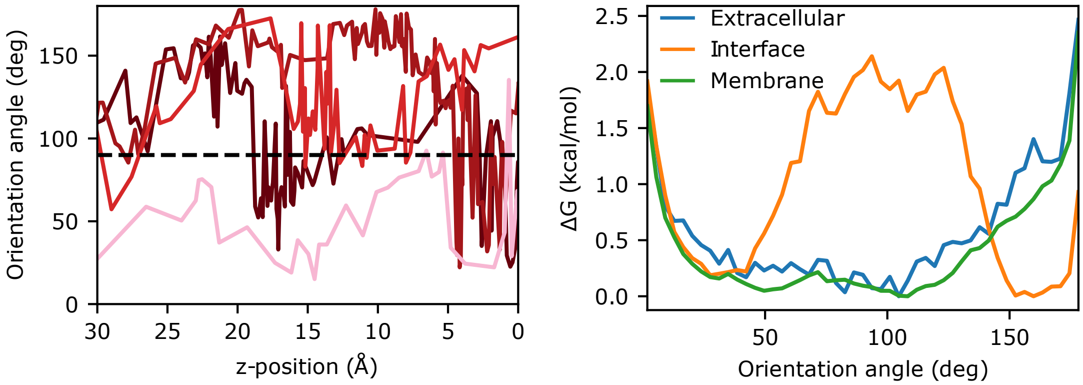


Figure S7: **Fentanyl prefers a vertical orientation at the interface in simulation set 1. Left.** Fentanyl orientation angle along stitched histories of walkers reaching the center of the membrane. **Right.** Potential mean force along the orientation angle while fentanyl is in the extracellular region, at the membrane interface, and within the membrane.

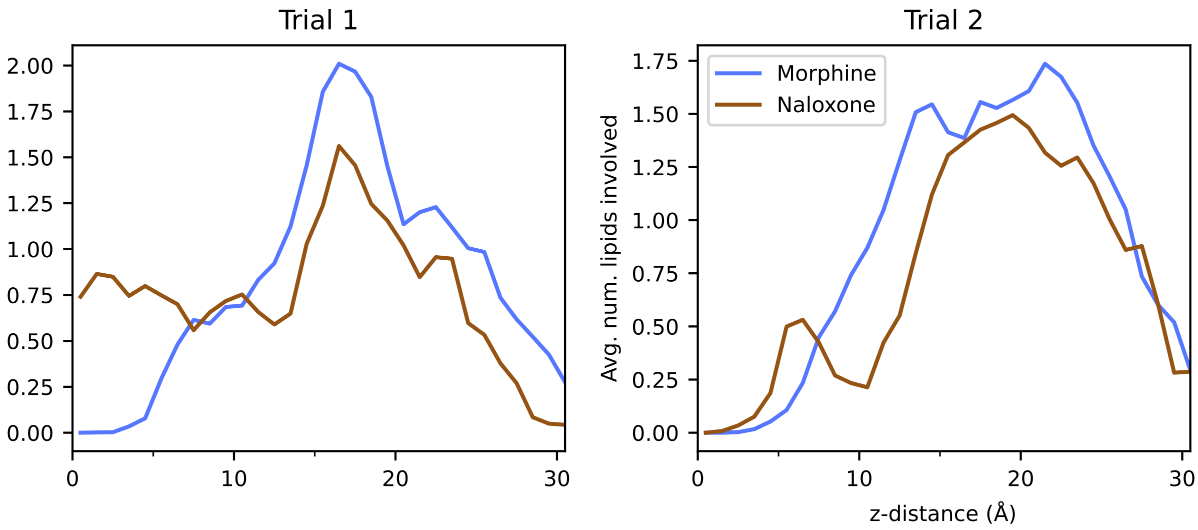


Figure S8: **Morphine and naloxone can interact with two separate phospholipid molecules while at the membrane interface.** Average number of lipids involved in hydrogen bonds with either morphine or naloxone, as a function of z-distance.

A

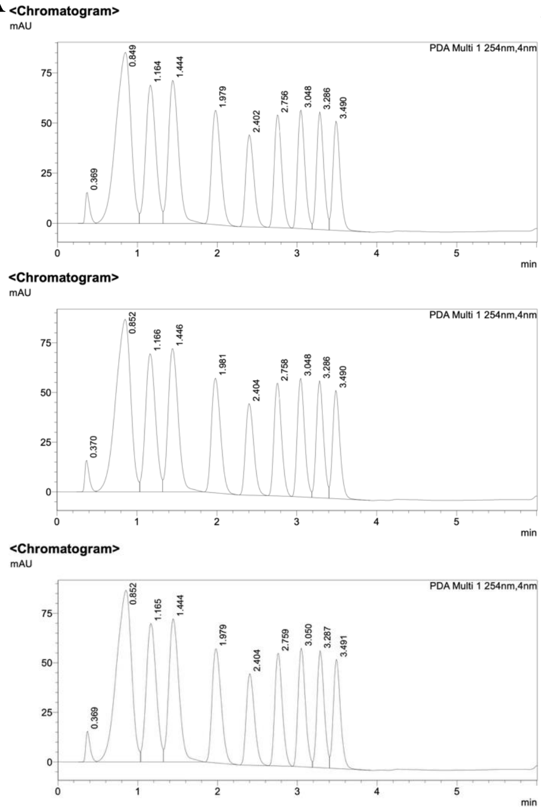


B

| Calibration Standards | Structure | CHI (IAM)[54] | $t_R \pm$ SD (mins) n=8 |
|---|---|---|---|
| octanophenone | 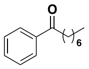 | 49.4 | 3.48 ± 0.04 |
| heptanophenone | 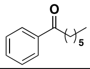 | 45.7 | 3.28 ± 0.04 |
| hexanophenone | 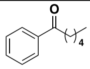 | 41.8 | 3.04 ± 0.04 |
| valerophenone | 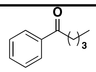 | 37.3 | 2.75 ± 0.04 |
| butyrophenone | 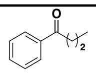 | 32.0 | 2.39 ± 0.04 |
| propiophenone | 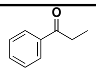 | 25.9 | 1.97 ± 0.04 |
| acetophenone | 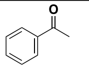 | 17.2 | 1.44 ± 0.04 |
| acetanilide | 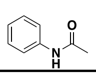 | 11.5 | 1.16 ± 0.05 |
| paracetamol | 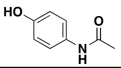 | 2.90 | 0.84 ± 0.05 |

Figure S9: a) Typical chromatograms for the IAM calibration mixture generated using method 1 with UV detection at 254 nm. b) Table of IAM calibration mixture components, their literature CHI(IAM)[S5] and average $t_R$±SD from n=8 repeats.

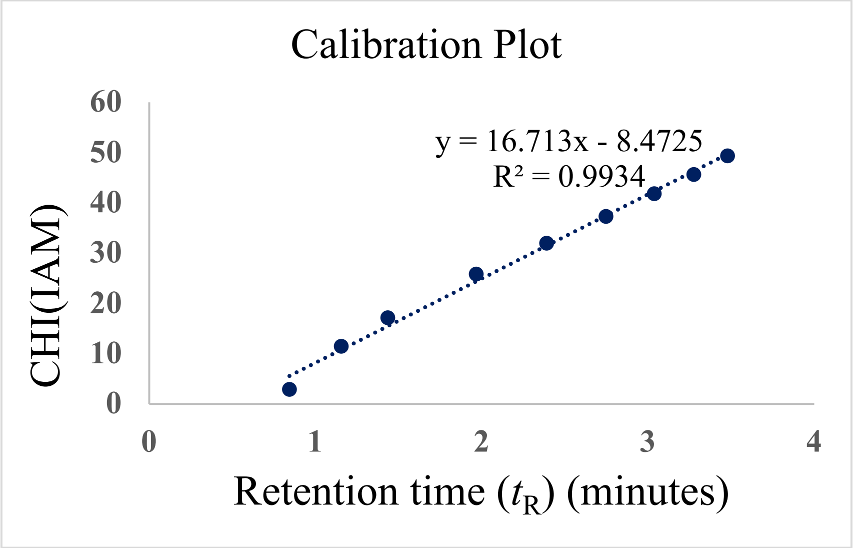


Figure S10: Calibration plot showing literature CHI (IAM)[S5] values against the $t_R$ measured for each component in the IAM calibration mixture. The data is fitted with a straight-line y = 16.713x - 8.4725, correlation coefficient ($r^2$) $>$ 0.99.

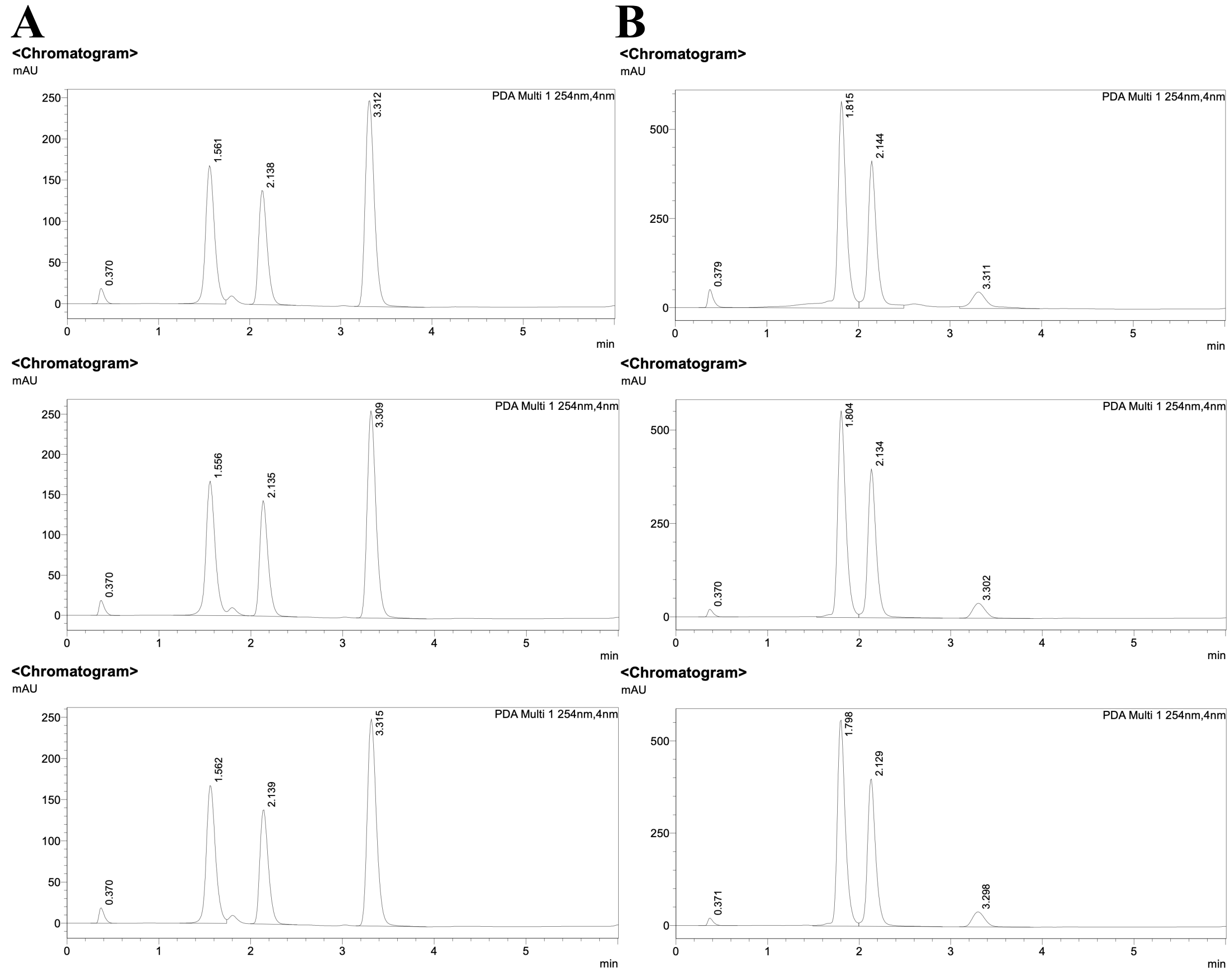


Figure S11: a) Typical chromatograms for suitability Test mixture 1 (nicardipine, warfarin and carbamazepine) and b) 2 (propanolol, indomethacin and colchicine) generated using method 1 with UV detection at 254 nm. All suitability tests were carried out daily in triplicate and the data summarized in Table S4.

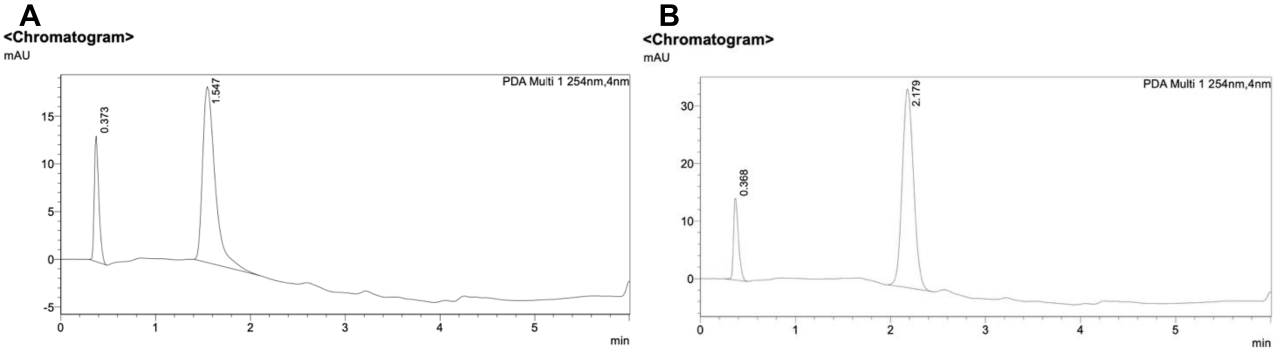


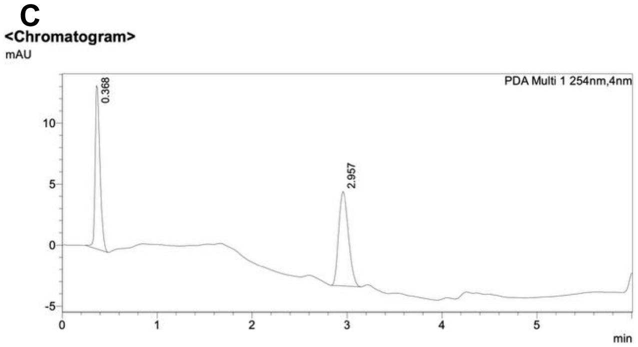


**D**

| Molecule | $t_R$ ± SD (mins) n=6 | CHI(IAM) |
|---|---|---|
| Morphine | 1.55 ± 0.03 | 17 |
| Naloxone | 2.18 ± 0.05 | 28 |
| Fentanyl | 2.96 ± 0.05 | 41 |

Figure S12: Typical chromatograms for a) morphine; b) naloxone; c) fentanyl generated using method 1 with UV detection at 254 nm. d) The mean $t_R$±SD of each molecule from n=6 repeats and the corresponding CHI(IAM) determined using calibration plot (Figure S9) is shown.

(S2) Alexander, S. P. H.; Christopoulos, A.; Davenport, A. P.; Kelly, E.; Mathie, A. A.; Peters, J. A.; Veale, E. L.; Armstrong, J. F.; Faccenda, E.; Harding, S. D.; Davies, J. A.; Abbracchio, M. P.; Abraham, G.; Agoulnik, A.; Alexander, W.; Alhosaini, K.; Bäck, M.; Baker, J. G.; Barnes, N. M.; Bathgate, R.; Beaulieu, J.-M.; Beck-Sickinger, A. G.; Behrens, M.; Bernstein, K. E.; Bettler, B.; Birdsall, N. J. M.; Blaho, V.; Boulay, F.; Bousquet, C.; Bräuner-Osborne, H.; Burnstock, G.; Caló, G.; Castaño, J. P.; Catt, K. J.; Ceruti, S.; Chazot, P.; Chiang, N.; Chini, B.; Chun, J.; Cianciulli, A.; Civelli, O.; Clapp, L. H.; Couture, R.; Cox, H. M.; Csaba, Z.; Dahlgren, C.; Dent, G.; Douglas, S. D.; Dournaud, P.; Eguchi, S.; Escher, E.; Filardo, E. J.; Fong, T.; Fumagalli, M.; Gainetdinov, R. R.; Garelja, M. L.; de Gasparo, M.; Gerard, C.; Gershengorn, M.; Gobeil, F.; Goodfriend, T. L.; Goudet, C.; Grätz, L.; Gregory, K. J.; Gundlach, A. L.; Hamann, J.; Hanson, J.; Hauger, R. L.; Hay, D. L.; Heinemann, A.; Herr, D.; Hollenberg, M. D.; Holliday, N. D.; Horiuchi, M.; Hoyer, D.; Hunyady, L.; Husain, A.; IJzerman, A. P.; Inagami, T.; Jacobson, K. A.; Jensen, R. T.; Jockers, R.; Jonnalagadda, D.; Karnik, S.; Kaupmann, K.; Kemp, J.; Kennedy, C.; Kihara, Y.; Kitazawa, T.; Kozielewicz, P.; Kreienkamp, H.-J.; Kukkonen, J. P.; Langenhan, T.; Larhammar, D.; Leach, K.; Lecca, D.; Lee, J. D.; Leeman, S. E.; Leprince, J.; Li, X. X. et al. The Concise Guide to PHARMACOLOGY 2023/24: G Protein-Coupled Receptors. *Br. J. Pharmacol.* **2023**, *180*, S23–S144.

(S3) Willighagen, E. L.; Mayfield, J. W.; Alvarsson, J.; Berg, A.; Carlsson, L.; Jeliazkova, N.; Kuhn, S.; Pluskal, T.; Rojas-Chertó, M.; Spjuth, O.; Torrance, G.; Evelo, C. T.; Guha, R.; Steinbeck, C. The Chemistry Development Kit (CDK) v2.0:

Atom Typing, Depiction, Molecular Formulas, and Substructure Searching. *J. Cheminform.* **2017**, *9*, 33.

(S4) Zhang, S.; Thompson, J. P.; Xia, J.; Bogetti, A. T.; York, F.; Skillman, A. G.; Chong, L. T.; LeBard, D. N. Mechanistic Insights into Passive Membrane Permeability of Drug-like Molecules from a Weighted Ensemble of Trajectories. *J. Chem. Inf. Model.* **2022**, *62*, 1891–1904.

(S5) Valko, K.; Du, C. M.; Bevan, C. D.; Reynolds, D. P.; Abraham, M. H. Rapid-Gradient HPLC Method for Measuring Drug Interactions with Immobilized Artificial Membrane: Comparison with Other Lipophilicity Measures. *J. Pharm. Sci.* **2000**, *89*, 1085–1096.
